# Evaluating the capabilities of the Astral mass analyzer for single-cell proteomics

**DOI:** 10.1101/2023.06.06.543943

**Authors:** Valdemaras Petrosius, Pedro Aragon-Fernandez, Tabiwang N. Arrey, Nil Üresin, Benjamin Furtwängler, Hamish Stewart, Eduard Denisov, Johannes Petzoldt, Amelia C. Peterson, Christian Hock, Eugen Damoc, Alexander Makarov, Vlad Zabrouskov, Bo T. Porse, Erwin M. Schoof

## Abstract

The complexity of human physiology arises from well-orchestrated interactions between trillions of single cells in the body. While single-cell RNA sequencing (scRNA-seq) has enhanced our understanding of cell diversity, gene expression alone does not fully characterize cell phenotypes. Additional molecular dimensions, such as proteins, are needed to define cellular states accurately. Mass spectrometry (MS)-based proteomics has emerged as a powerful tool for comprehensive protein analysis, including single-cell applications. However, challenges remain in terms of throughput and proteomic depth, in order to maximize the biological impact of single-cell proteomics by Mass Spectrometry (scp-MS) workflows. This study leverages a novel high-resolution, accurate mass (HRAM) instrument platform, consisting of both an Orbitrap and an innovative HRAM Asymmetric Track Lossless (Astral) analyzer. The Astral analyzer offers high sensitivity and resolution through lossless ion transfer and a unique flight track design. We evaluate the performance of the Thermo Scientific Orbitrap Astral MS using Data-Independent Acquisition (DIA) and assess proteome depth and quantitative precision for ultra-low input samples. Optimal DIA method parameters for single-cell proteomics are identified, and we demonstrate the ability of the instrument to study cell cycle dynamics in Human Embryonic Kidney (HEK293) cells, and cancer cell heterogeneity in a primary Acute Myeloid Leukemia (AML) culture model.

## 1. Introduction

The multicellular intricacy governing human physiology is the result of a landscape of coordinated interactions between trillions of single cells that constitute the human body. With the arrival of single-cell RNA sequencing (scRNA-seq), our understanding about the phenotypic diversity present in cell populations previously thought to be of discrete nature has rapidly increased^1–3^. However, gene expression on its own does not capture the complete context required to characterize the phenotypic state of the cell^4–7^. Thus, modalities expanding across other dimensions of molecular information, such as proteins, are necessary to gain fundamental understanding on how different cellular states are truly defined.

In the past, measuring protein abundance levels at single-cell resolution was predominantly restricted to methods that utilized affinity or chemical reagents to initially label a protein of interest to then estimate its abundance by using techniques such as immunofluorescence microscopy, fluorescence activated cell sorting (FACS) or next-generation sequencing (NGS) in conjunction with oligonucleotide-linked reagents^8, 9^. In spite of the wide availability of methods, the direct and systemic quantification of proteins, both at a global scale and at the single-cell level remains challenging.

Over the last three decades, mass spectrometry (MS)- based proteomics has established itself as a powerful analytical tool for the comprehensive characterization of proteins contained in a biological sample. However, only since recent advancements enhancing all aspects of the analytical framework of MS-based proteomics, ranging from sample preparation to data processing^10–15^, have allowed the field to start branching out into single-cell applications.

Although single-cell proteomics by mass spectrometry (scp-MS) is capable of quantifying 1000-2000 protein groups at a moderately applicable throughput^11, 16–19^, it is still subject to significant limitations that need to be addressed. Among these, key improvements lie in the number of cells that can be analyzed per unit time and the proteomic depth that can be accurately reached. Studies employing scp-MS have mostly been conducted using Orbitrap (OT) and Time of Flight (TOF) analyzers^16, 18, 20, 21^, however, the implementation of Linear Ion Traps (LIT) for this purpose has recently been successfully demonstrated by us and others^22–24^.

Integrating alternative mass analyzers such as LIT highlighted the advantages that highly sensitive mass analyzers have for scp-MS, albeit that the advantages of LIT are limited by its inherently low resolution^23, 25^. A principal obstacle is posed by the need to quantify the miniscule amount of ions that can be produced from a single cell, thereby putting sensitivity at the very forefront of priorities in terms of mass analyzer characteristics. While resolution of OT-based data acquisition remains unsurpassed, extensive ion injection times (IIT) are required, ranging in the hundreds of milliseconds for identification and accurate quantification of single-cell signal^18, 26–28^, underlining the need for mass analyzers that strike an optimal balance between sensitivity and high-resolution resolving power.

Here, we leverage a novel high-resolution accurate mass instrument platform (Figure 1A) containing an Orbitrap and a newly developed high-resolution/accurate mass (HRAM) Asymmetric Track Lossless (Astral) analyzer to study biological variation in different model cell systems^29^. The Astral is a novel mass analyzer that shares some of the operational principles with established mass analyzers, including the Orbitrap, ion trap and time-of-flight. However, its nearly lossless ion transfer and therefore high sensitivity is its most unique and consequential property^30^. Additionally, the asymmetric flight track of over 30 meters allows the Astral analyzer to achieve high resolution (80,000 at m/z 524) without compromising ion transmission at scan speeds up to 200 Hz. In this study, we evaluate the performance of the Thermo Scientific^TM^ Orbitrap^TM^ Astral^TM^ Orbitrap Astral MS using Data-Independent Acquisition (DIA) and assess the proteome depth and quantitative precision of the obtained data for ultra-low input samples. Furthermore, we carry out survey experiments to identify optimal DIA method parameters for scp-MS application. Finally, we apply our established method to study the biological variation present at the protein-level in two model systems: 1) cell cycle in Human Embryonic Kidney (HEK293) cells, and 2) cancer cell heterogeneity in a primary Acute Myeloid Leukemia (AML) culture model.

**Figure 1.**
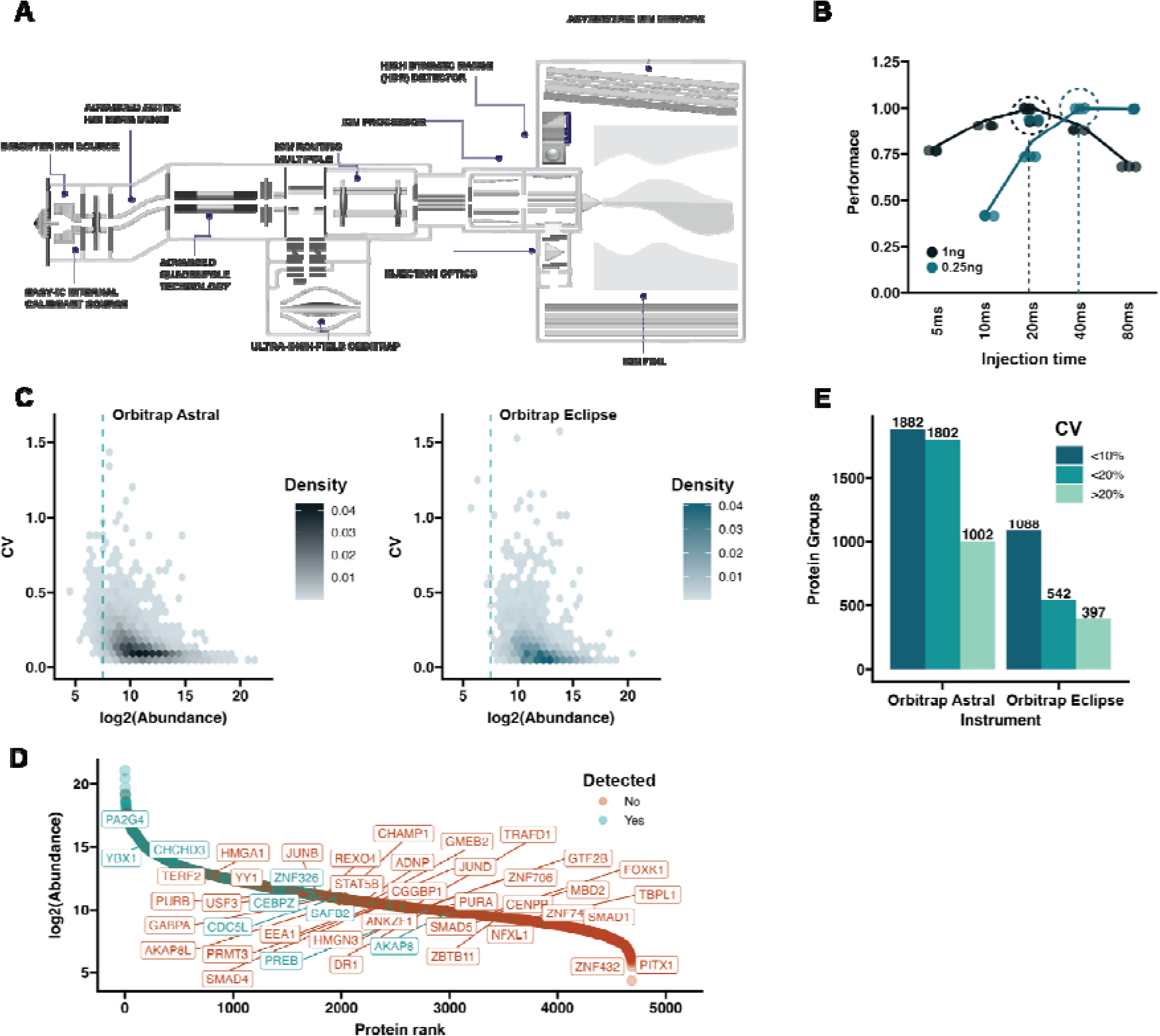
Optimizing Astral parameters for limited-input proteomics. **A)** Orbitrap Astral MS instrument schematic. Three main components constitute the analytical architecture of the Orbitrap Astral MS: i) a quadrupole mass filter, ii) an Orbitrap analyzer and iii) the newly developed HRAM Astral analyzer for fast parallelized spectral acquisition of fragment ions. **B)** Protein groups identified from 1ng (black) and 250pg (turquoise) of Pierce HeLa digest, with different MS2 injection times on the Astral analyzer. **C)** Hexbin plots the coefficient of variation (CV) distribution with respect to the log2 transformer protein abundance. **D)** Rank plot based on Astral protein abundances, the proteins detected with OT/OT on Eclipse are indicated by different colors. Transcription factors are labeled with their respective gene names **E)** Barplots showing the total number of proteins identified below a certain CV threshold with Orbitrap Astral or Orbitrap Eclipse instruments from 1ng. Detected transcription factors highlighted.

## 2. Results

### The Astral analyzer for limited input proteomics

To maximize the proteome coverage we adopted our recently established, limited input tailored, wide isolation window HRMS1 (WISH) - DIA method, which is focused on MS1 level quantification and uses MS2 information only for identification purposes^21^. Due to the novelty of the Astral mass analyzer, the optimal instrument parameters used for ultra-low input samples were unknown. To systematically investigate key parameters such as IIT we first used a HeLa dilution series to evaluate appropriate IIT to maximize proteome coverage without sacrificing quantitative precision. The high sensitivity of the Astral analyzer, combined with its high scan rate should alleviate the burden of long IIT required for ultra-low input samples. Therefore, we hypothesized that we could in turn utilize narrower isolation windows compared to OT-platforms, generating less complex fragment spectra and increasing spectral specificity. To test this, we designed a panel of acquisition methods using a combination of OT and Astral, where the precursor spectra were measured at a constant resolution of 120,000 on the OT, while varying the maximum fragment IIT into the Astral analyzer (see Methods). The differences in scan cycle time were compensated by concomitantly widening the isolation windows. The length of the total scan cycle time was coordinated with the chosen liquid chromatography (LC) method, utilizing the uPAC Neo “Low Load” analytical column. Using a total method time of 18 minutes sample-to-sample, we can process 80 samples per day (Figure S1A), while having a mean peak full-width at half maximum (FWHM) of 2.8 seconds (Figure S1B-C). The chosen cycle time of 0.7 seconds ensures that enough data points are collected per peptide elution peak to ensure accurate chromatographic peak approximation (Figure S1E-F).

We applied our designed OT/Astral methods on two injection loads (1ng and 250pg) of Pierce Hela digest to determine the parameters that would yield the deepest proteome profile (Figure 1B). In line with the enhanced sensitivity of the Astral analyzer, the highest identification number was already achieved with 20ms injection time, compared to 246ms on a previous OT-only platform^21^. Decreasing the injected peptide amount to 250pg shifted the optimal injection time to 40ms (Figure 1B), recapitulating our previous finding that maximal proteome coverage for ultra-low input samples is obtained with long IIT and large isolation windows, albeit with substantially shorter IIT than an OT-only platform^21^. To directly compare the quantitative precision and proteome coverage of the Orbitrap Astral mass spectrometer we analyzed 1ng Hela digest on an Orbitrap

Eclipse mass spectrometer with an optimized OT/OT WISH-DIA method and used that as a reference point. We could clearly see that the OT/Astral WISH-DIA method allows us to quantify proteins that have a lower abundance compared to OT/OT (Figure 1C-D). Astonishingly, leveraging the Astral analyzer not only doubled the total number of identified proteins, but also maintained a reminiscent degree or precision, where the majority of proteins fell below a CV value of 20% (Figure 1E). This becomes particularly significant, when key cellular decision driving molecules, such as transcription factors are of interest. The high sensitivity of the analyzer allowed us to quantify a remarkably higher number of transcription factors that otherwise remain hidden, when analysis is carried from such miniscule amount of sample (Figure 3D). Overall, our data underlines the new Astral analyzer to exhibit enhanced capacity to facilitate deep proteome profiling, while maintaining quantitative precision.

### The Orbitrap Astral MS for single-cell proteomics

With the initial parameter boundaries for ultra-low input established, we extended our optimization to cover actual single-cell input. We prepared single-cell samples with a standard FACS workflow in a 384-well plate format (see Methods^18, 19^) and generated an additional series of acquisition methods to pinpoint the optimal parameters for scp-MS (see Methods). In line with the increased sensitivity of the Astral analyzer, the best performance was observed with 65ms IT (Figure 2A), which is a considerable decrease compared to 246-504ms required for OT methods^18, 26–28^. Moreover, we managed to identify ∼2000 protein groups, underlining superb achievable proteome depth at much higher throughput relative to previous label-free scp-MS methods^16, 31^.

**Figure 2.**
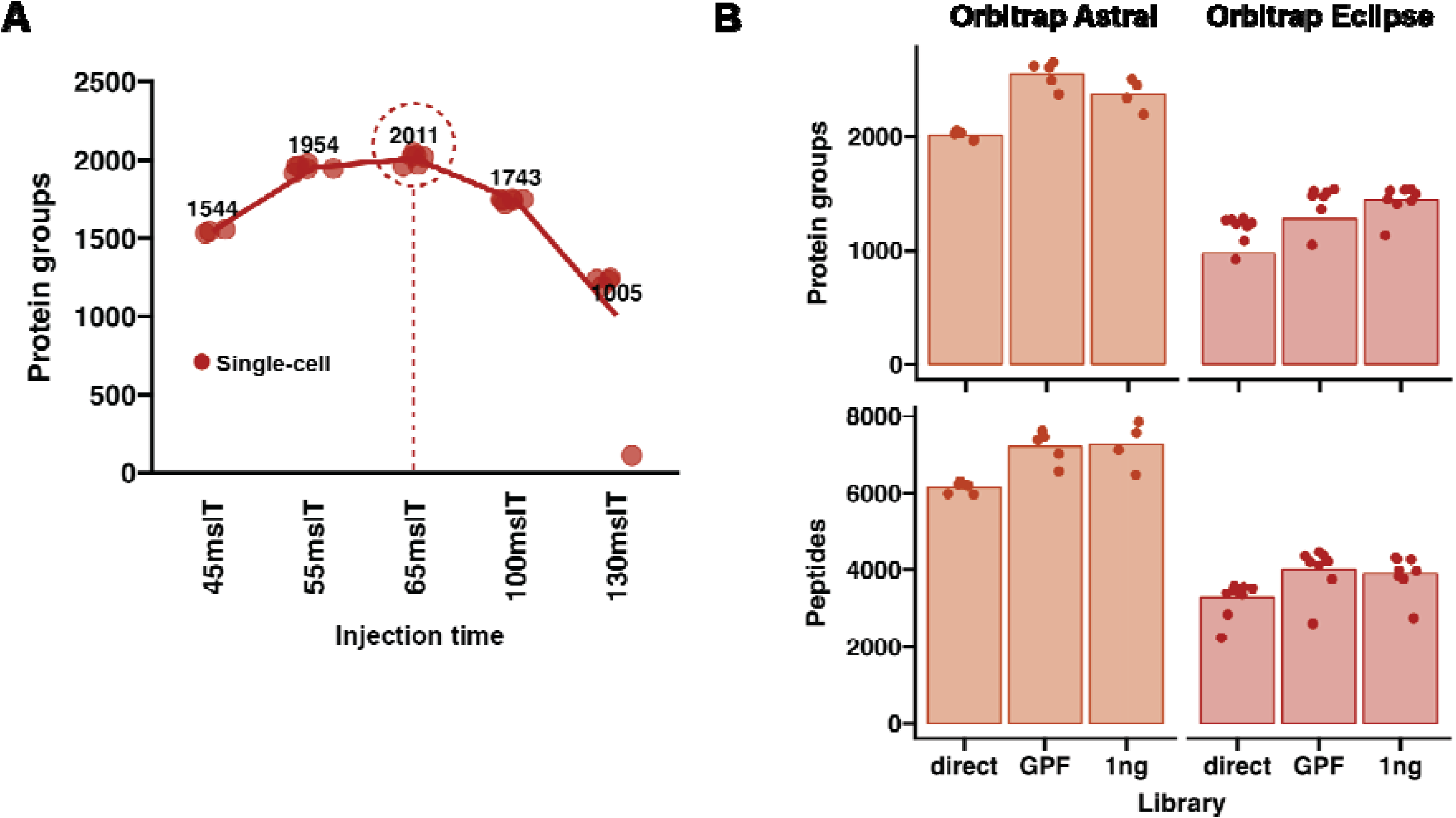
Performance of Orbitrap Astral MS for single-cell proteome profiling. **A)** Method optimization for actual single-cell samples **B)** Comparison of OT/Astral and OT/OT methods (Orbitrap Eclipse Tribrid) with different DIA search strategies.

Comparing the performance of scp-MS workflows across instrument platforms is challenged by the wide range of variables inherent to the experimental workflow. Main contributors to this variation are cells of interest, sample preparation, LC separation, gradient lengths and type of ion mobility^16, 31–33^. Accordingly, to gauge the enhanced capacity of the Astral analyzer for scp-MS compared to previous platforms in our lab, we analyzed and compared the same type of single-cell samples, run under near-identical LC conditions, with an OT/OT method on an Orbitrap Eclipse Instrument (Figure 2B). Here, leveraging the Astral doubled the number of identified proteins and peptides (Figure 2A). To push the proteome depth limits further, we evaluated the efficiency of Gas-Phase Fractionation (GPF, ^34^) and 1ng Hela digest libraries, and observed their ability to further boost identifications by ∼20%, reaching close to ∼2500 protein groups per single-cell (Figure 2A). Overall, these results highlight the ability of the Orbitrap Astral MS to augment the application of scp-MS, and substantially alleviate some of the constraints of having to compromise cell throughput and proteome coverage.

### Exploring biological variation during the cell cycle

As the Orbitrap Astral MS exhibited superb proteome coverage from single-cell input, we leveraged our optimized method to generate a small ∼100 cell data set to see if the resulting data allows us to study core cellular processes such as the cell cycle. In this larger-scale dataset, we obtained ∼2400 protein groups per single-cell and a reasonably high level of data completeness at the protein-level wise (Figure 3A-B). To determine if the data captures latent features that govern the cell cycle progression, we used two dimensional reduction techniques (PCA and UMAP) to account for both linear and non-linear trends in the data (Figure 3C). Intriguingly, the cells assumed a quasi-circle like pattern, which would be expected for a cyclical process as the cell cycle. The UMAP also resembles a cycling pattern, albeit to a lesser extent. Next, to gain insight into which section corresponds to specific cell cycle stages, we mapped protein expression levels of two classical cell cycle proteins: 1) Geminin (GMNN), which peaks in G2/M cell cycle stages and 2) PCNA, which is a well known S-phase marker (refs), on to the PCA and UMAP (Figure 3). The expression pattern of the proteins showed distinct patterns and had highest intensity in specific sections of the PCA and UMAP, indicating those specific areas to contain G2/M and S-phase like cells. To boost the confidence in the identified cell cycle stages, we used known G1- and G2- specific proteins (?) to calculate an aggregated score for these cell cycle stages(refs). Cells that possessed an overall lower value of PC1 had a higher G2 score, whereas the opposite was true for G1 scores. Furthermore, we observed cells with the highest levels of PCNA to be found between cells with high G1 and G2 scores, following the canonical cell cycle transition scheme (Figure 3C).

To simplify our exploration of the protein expression dynamics during the cell cycle, we capitalized on the circular nature of the data representation and utilized a geometric approach to convert the principal component (PC) values to a single angular value, acting as a singular interpretation of the cell cycle trajectory (Figure 3D). To ensure the robustness of this approach, we inspected the expression levels of one the key drivers of the cell cycle CDK1/CyclinB (CCNB1) complex. As expected, CDK1 levels remain rather constant throughout the cell cycle, while Cyclin B levels peak at a specific trajectory value (Figure 3D). Furthermore, we could see PCNA levels peak before CyclinB, in accordance with classical knowledge.

Another pivotal molecular engine is the MCM complex that is crucial for proper DNA replication and histone recycling, which if compromised, can have tremendous ramifications^35–37^. Expression of MCM components during the cell cycle has been studied with numerous techniques, including immunofluorescent imaging-based tools with inherent single-cell resolution^38^. However, our approach allows us to investigate these patterns in a data-driven manner, and without the biases introduced by affinity based reagents or endogenous fluorescently tagged protein expressions^39^. We plotted the circular protein abundance trends for the 6 MCM components quantified in our data (Figure 4E). A distinct trend for each subunit was observed, which is expected considering the complicated mechanism of MCM regulation^38^. Relative to PCNA, it seems that MCM expression is correlated with S-phase, although MCM3/4 subunit levels peak at a later stage (Figure 3E). Although further biological experiments would be required to conclusively and orthogonally test the validity of our observations, our data underlines the ability of Orbitrap Astral MS driven scp-MS workflows to generate biological hypotheses, requiring less than 2 days of LC-MS instrument time.

**Figure 3.**
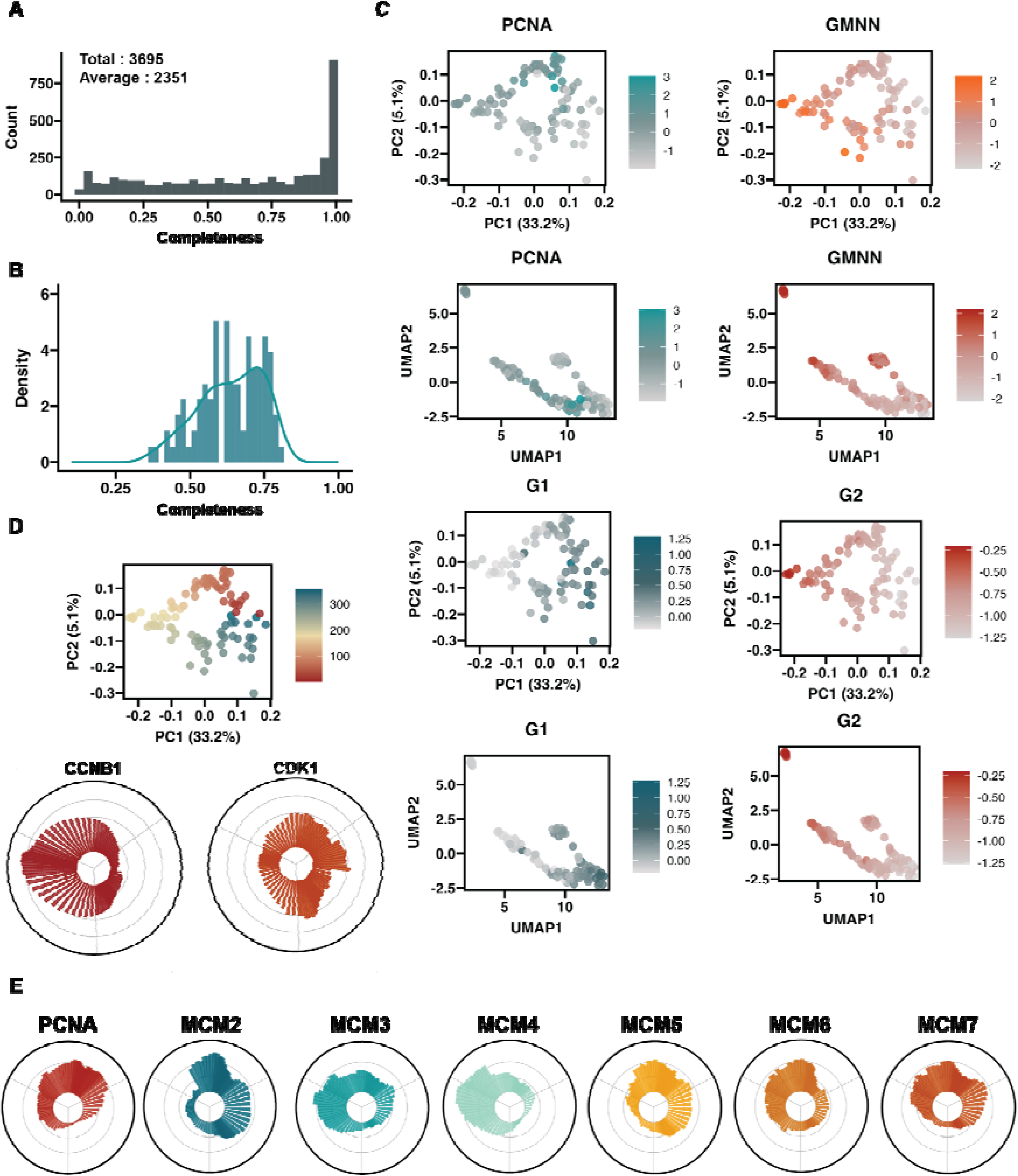
Exploring protein-derived biological variation in the cell cycle. **A)** Histogram of protein wise data completeness. **B)** Histogram of cell-centric data completeness **C)** Parametric (PCA) and non-parametric (UMAP) clustering analysis of the generated single-cell data. Different protein abundance values or cell cycle state scores are indicated by color intensities. **D)** Visualization of the geometrically infer cell cycle trajectory with circular plots showing expression of CCNB1 and CDK1. **E)** Circular plots of MCM complex subunit expression along the cell cycle trajectory.

**Figure 4.**
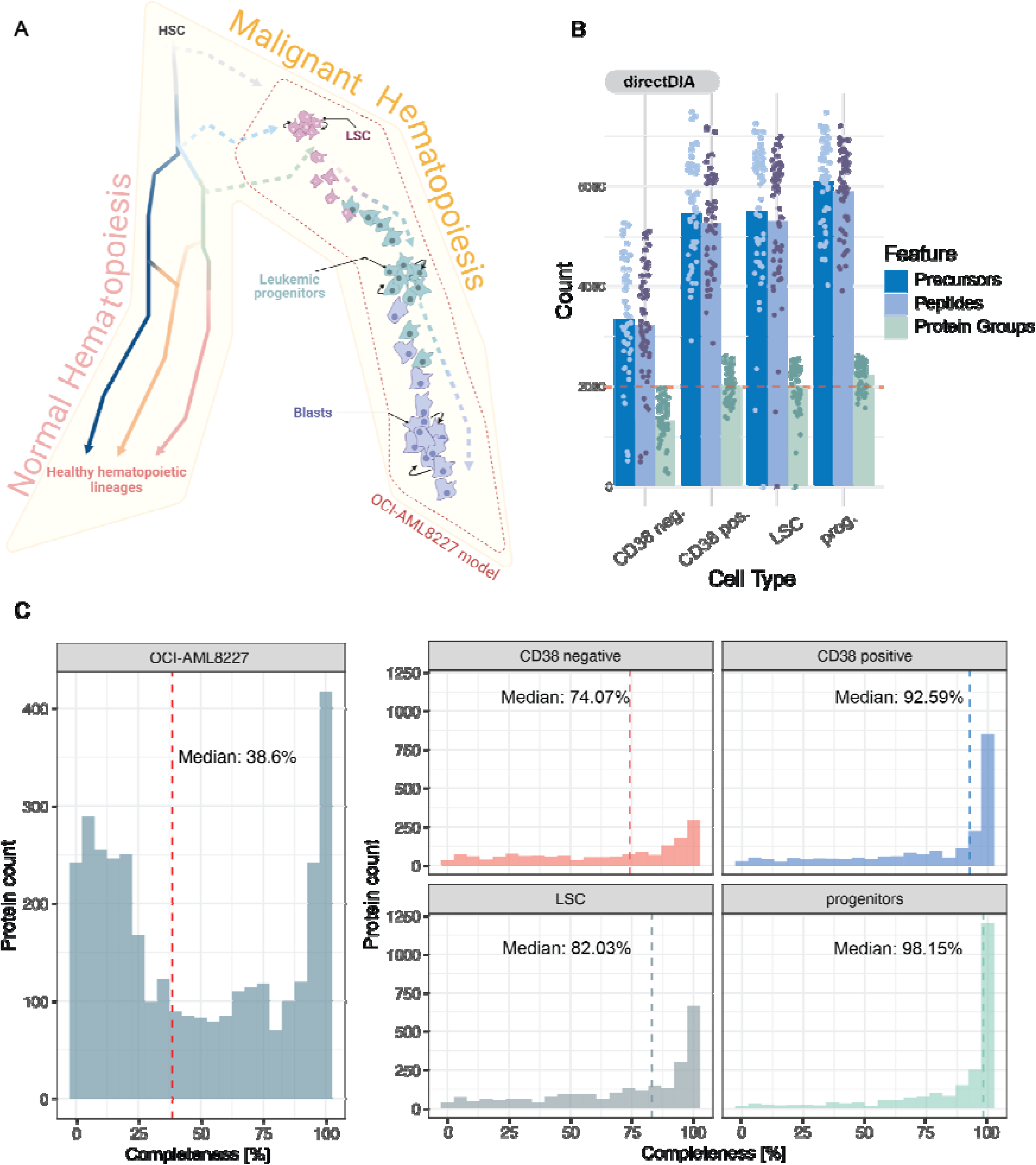
Single cell proteome analysis of the OCI-AML8227 hierarchy. **A)** Differentiation continuum during normal and abnormal hematopoiesis. Under normal circumstances, quiescent and self-renewing hematopoietic stem cells (HSCs) reside at the apex of the cellular hierarchy. From these stem cells, different lineage-committed cells that constitute the blood system emerge. Mutations across the healthy differentiation progression lead to the development of AML (dashed arrows), where self-renewing leukemic stem cells (LSCs) are located at the apex of the abnormal hierarchy and differentiate into leukemic progenitors and finally blasts. This malignant differentiation progression is recapitulated by the OCI-AML8227 model (highlighted in fine-dashed red). **B)** Library-free based identification summary of each cell population isolated from the OCI-AML8227 system. 216 cells were acquired in total. Each dot represents a single cell. **C)** Assessment of protein data completeness. Left panel accounts for the overall protein data completeness across all populations. Right panel represents the overall protein data completeness within each population. Red dashed lines display the median.

### Characterization of cellular heterogeneity in an acute myeloid leukemia hierarchy

To assess the capabilities of our analytical setup in a biologically relevant and heterogeneous setting, we selected the OCI-AML8227 system^40^, a primary Acute Myeloid Leukemia (AML) model that preserves the hierarchical nature of AML while cultured ex-vivo. The model comprises self-renewing leukemic stem cells (LSCs; CD34+, CD38-) positioned at the hierarchical apex. These LSCs undergo differentiation into progenitor cells (CD34+, CD38+) and subsequently transition into blasts, where their CD34 expression is lost. Blasts are characterized by the coexistence of both CD38+ and CD38- cell populations (Figure 4A). To assess the ability of the Orbitrap Astral MS to detect cell heterogeneity in highly similar cells derived from the same heterogeneous cell system, we acquired 54 single cells each, derived from four populations based on their CD34/CD38 marker expression and analyzed them with the WISH-DIA data acquisition method described above. Despite the OCI-AML8227 cells being derived from primary cells, rather than immortal cell lines such as HEK293 or HeLa, and as a consequence are expected to express fewer proteins, our analysis yielded the identification of 3892 protein groups across all cell populations. Furthermore, our findings consistently demonstrated an average identification of over 2000 protein groups per cell within each population, with the exception of the CD38 negative blasts, where the count was approximately 1300 protein groups (Figure 4B). Upon further exploration of the overall data quality, we observed the degree of protein data completeness to exhibit a median value of 38%. However, when considering each population individually, the degree of completeness increased significantly, ranging from a minimum median value of 74% to a maximum of 98% (Figure 4C). These results underline the extent of heterogeneity that exists within the OCI-AML8227 cell culture model, and that the heterogeneity can be decreased by enriching for specific cell-types through FACS-based means.

The favorable number of identifications observed motivated us to conduct a more comprehensive analysis of the acquired data and evaluate its possible biological relevance. Upon closer examination, we discovered a broad range of protein abundances, highlighting the dynamic range present in the data. Among these proteins, we identified several kinases, transcription regulators, and epigenetic regulators of interest (Supplementary tables 1). Notably, a large proportion of these proteins were predominantly ranked in the mid-to-lower end of the abundance distribution (Figure 5A). To get a better perspective of the gained proteomic depth, we compared our results with a dataset previously generated in-house from the same leukemia model using an Orbitrap Eclipse mass spectrometer^19^. We observed a close to 8-fold increase in the number of transcriptional regulators that we could identify, and a 2-fold gain for kinases. Similar numbers were obtained for epigenetic regulators. These results further emphasize the additional biological information that can be gained by scp-MS, when facilitated by the enhanced sensitivity of the Orbitrap Astral MS.

**Figure 5.**
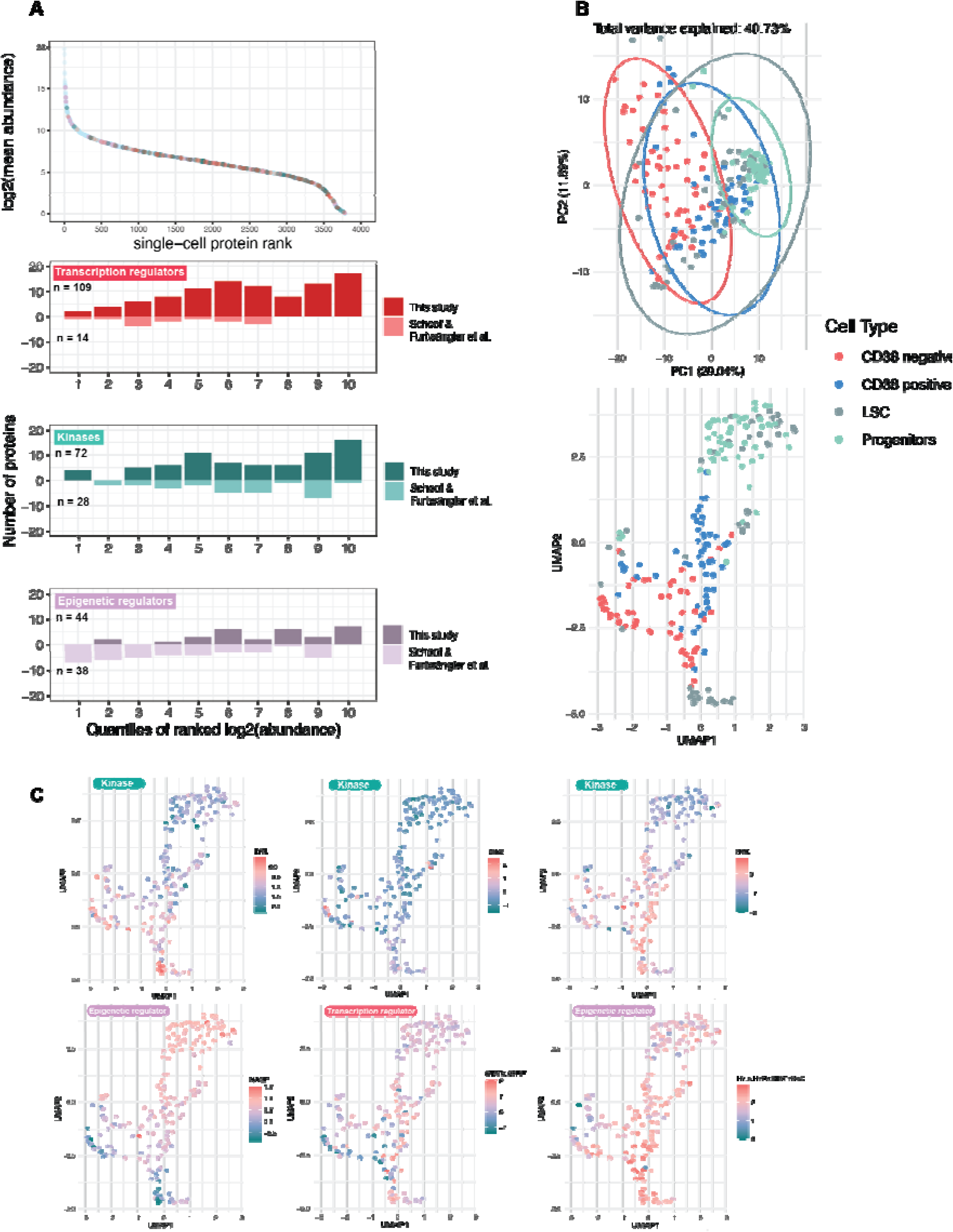
Biological examination of the cell heterogeneity present in the OCI-AML8227 model system. **A)** Upper panel: Rank plot of proteins identified across all populations present in the OCI-AML8227 model. Highlighted dots represent kinases, epigenetic-and transcription regulators of interest. Lower panel: Protein distribution and identification comparison between this study and data generated by Schoof & Furtwängler et al. **B)** Upper panel: Principal component analysis of all cell populations. Ellipses represent 95 % confidence intervals. Lower panel: UMAP embedding of all cells. **C)** UMAP embedding of all cells overlaid with the scpMS-derived protein expression of selected proteins.

Next, we sought to evaluate the capability of our analytical setup in detecting known heterogeneity within the OCI-AML8227 model system. Separation of all acquired cells by their principal components revealed the clear manifestation of phenotypic diversity within our leukemia model (Figure 5B). In particular, LSCs and progenitors exhibited lesser variance between each cell type, indicating their close resemblance. Conversely, the CD38 +/-blasts displayed a more scattered distribution within their respective populations, but still exhibited distinguishable characteristics from one another as well as from the other cell populations. This pattern reflects the presence of multiple differentiation stages across the cellular hierarchy, which are better captured by applying non-linear methods. The generated UMAP embedding resulted in a clear recapitulation of the known differentiation trajectories of OCI-AML8227, starting with LSCs that progress into progenitors and finally convert into CD38 negative blasts. Remarkably, these results demonstrated a notable proportion of LSCs bearing a resemblance to the CD38 negative population. These findings align with previous observations made by our group, indicating that LSCs have the potential to directly differentiate into CD38 negative blasts^19^ (Figure 5B). Notably, our findings revealed that many differentially expressed (DE) proteins exhibited heterogeneous expression patterns across the various cell differentiation stages. This suggests that our single-cell data effectively captures gradual changes in protein expression that would typically remain undetected in bulk data (Figure 5C).

## Discussion

In this study, we benchmark the potential application of the novel Thermo Fisher Orbitrap Astral MS for studying biological variation at single-cell resolution. Given the nascent nature of this instrument, we first needed to identify optimal data acquisition parameters for profiling single-cell proteomes. To narrow down the parameter search space and gain a better understanding of the capacity of the instrument, we first evaluated the IIT required for maximal proteome coverage from peptide digests diluted to ultra-low input (1ng) (Figure 1). Remarkably, with the established optimal parameters, incorporation of the Astral analyzer for fragment spectra acquisition doubled the proteome coverage comparing to previous generation MS instruments, unlocking numerous low abundant key proteins such as transcription factors (Figure 1C-D) with relatively high precision (Figure 1E). The resulting optimal method for ultra-low input utilized injection times that were almost an order of magnitude lower compared to the Orbitrap Eclipse instrument, underlining a marked increase in sensitivity, which is essential when profiling single cells^18, 26–28^. Similarly, we also found that for actual single-cell samples, the optimal method utilized an injection time that was markedly reduced (Figure 2A). Finally, the resulting proteome coverage represented double the number of proteins and peptides compared to previous OT/OT based methods (Figure 2B). Overall, these results substantiate the potential of the Orbitrap Astral MS to propel the field of scp-MS forward to new horizons.

To truly gauge the capacity of the Astral mass analyzer for capturing biological variation at single-cell resolution, and move beyond a purely technical evaluation, we used two biological systems possessing different degrees of complexity. The HEK293 system presents a rather homogenous population, for which the major variation present in the data is expected to be driven by cell cycle transition (Figure 3). Recently, we showed that deploying an asymmetric spectral library-driven search approach, using higher loads than the sample type of interest, can have negative effects on the generated data quality from low input samples^21^. To ensure maximum accuracy of the quantification of our single-cell data, and provide an unbiased evaluation of the sensitivity of our method, we here used a library-free approach to search our generated single-cell spectra (see Methods). We used dimensionality reduction techniques to analyse the global trends present in the data in both linear and non-linear space (Figure 3C). Strikingly, for the HEK293 results, we observed a clear circular pattern in the data in the PCA analysis. Upon examining the trends of canonical cell cycle regulated proteins, we observed a clear correlation along the principal components, underlining that we are capturing the latent protein-level variables driving the phenotypic variation in the data. We then use a geometric approach to infer a cell cycle trajectory and visualize the expression of the MCM helicase complex subunits throughout the cell cycle.

Finally, to assess the ability of Astral-driven scp-MS to interrogate cell heterogeneity at the protein level, we use the OCI-AML8827 primary cell model, which recapitulates the cell hierarchy found in AML. The resulting data revealed additional latent variables that govern the biological variation in this cellular system. Combined, our ultra-low input results suggest that the unique properties of the Astral analyzer allows us to capture a substantially larger number of crucial factors involved in cellular decision making (Figure 5A). The highly multi-dimensional nature of such scp-MS data precluded any clear separation of all the populations present in linear space (PCA). However, in non-linear space (UMAP) we could clearly see the ability of our proteomics data to separate the four key cell populations contained in OCI-AML8227, especially for the more distinct cell types such as blasts vs. progenitor/LSC. Moreover, we identified transcriptional regulators and kinases that are differentially expressed between the populations and showed that we are able to track their trends based on the cellular state/type. Compared to our previous TMTPro-based interrogation of this primary cell system, we drastically increased our ability to quantify these two protein classes in single cells (Figure 5A, 8-fold and 2.5-fold increase respectively). Resulting, this expansion of the scope of quantified key factors, we can obtain a more comprehensive understanding of the molecular intricacies governing malignant cell hierarchies. When taken together, this work underlines the impact of the increased sensitivity and speed of the Astral mass analyzer on our ability to advance our understanding of cell state heterogeneity at the proteome level.

## 3. Methods

### Cell culture and FACS sorting

HEK cells were cultured in RPMI media containing 10 % FBS and 1 % Penstrep. Upon 80% confluence, cells were harvested and washed three times with ice-cold PBS to remove any remaining growth media prior to cell sorting. The cells were resuspended in ice-cold PBS at 1e6 cells/ml prior to sorting. To sustain the hierarchical nature present in the OCI-AML8227 system, cells were grown in StemSpan SFEM II media, supplemented with growth factors (Miltenyi Biotec, IL-3, IL-6 and G-CSF (10 ng/mL), h-SCF and FLt3-L (50 ng/mL), and TPO (25 ng/mL). On the 6th day, cells were harvested, washed, counted, and resuspended in fresh StemSpan SFEM II media while constantly maintained on ice at a cell density of ∼5e6 cells/ml. Immunostaining was done for 30 mins on ice, using a CD34 antibody (CD34-APC-Cy7, Biolegend, clone 581) at 1:100 (vol/vol) and CD38 antibody (CD38-PE, BD, clone HB7) at 1:50 (vol/vol). Cells were washed with extra StemSpan SFEM II media, and subsequently underwent three washes with ice cold PBS to remove any remaining growth factors or other contaminants from the growth media.

Cell sorting was done on a FACS Aria III or Aria II instrument, controlled by the DIVA software package (v.8.0.2) and operated with a 100 μm nozzle. All cells were sorted at single-cell resolution, into a 384-well Eppendorf LoBind PCR plate (Eppendorf AG) containing 1 μL of lysis buffer (100 mM Triethylammonium bicarbonate (TEAB) pH 8.5, 20 % (v/v) 2,2,2-Trifluoroethanol (TFE)). Directly after sorting, plates were briefly spun, snap-frozen on dry ice for 5 min and then heated at 95 °C in a PCR machine (Applied Biosystems Veriti 384-well) for an additional 5 mins. Samples were then either subjected to further sample preparation or stored at −80 °C until further processing.

### Preparation of single cells for mass spectrometry

Well plates containing single-cell protein lysates were digested with 2 ng of Trypsin (Sigma cat. nr. T6567) supplied in 1 μL of digestion buffer (100 mM TEAB pH 8.5). The digestion was carried out overnight at 37 °C, and subsequently stopped by the addition of 1 μL 1 % (v/v) trifluoroacetic acid (TFA). The resulting peptides were either directly submitted to mass spectrometry analysis or stored at −80 °C until further processing. All reagent dispensing was done using an I-DOT One instrument (Dispendix).

### Liquid chromatography

Chromatographic separation of peptides was conducted on a vanquish Neo UHPLC system connected to a 50 cm uPAC Neo Low-load and an EASY-spray emitter via built-in NanoViper fittings (all ThermoScientific). All separations were carried out using a single-column configuration and with the column oven set to 50 °C. For single-cell derived peptides, the autosampler and injection valves were configured to perform direct injections from a 384 well plate using a 25 uL injection loop. 11.8 min gradients were applied where the percentage of buffer B (80 % ACN in H2O, 0.1 % FA) was initially increased from 0 to 13.5 % (0-2 min) with the nominal flow set at 750 nl/min. This was followed by gradual increase of buffer B from 13.6 to 99 % (2-5.1 min) at a flow rate of 200 nl/min and kept constant for 6.7 min (5.1-11.8).

### Mass spectrometry data acquisition

Acquisition of single-cell derived peptides was conducted with an Orbitrap Astral mass spectrometer operated in positive mode with the FAIMS Pro interface (Thermo Fisher Scientific) using a compensation voltage set to −50 V. Orbitrap MS^1^ spectra were acquired with the Orbitrap at a resolution of 120,000 and a scan range of 400 to 900 m/z with normalized automatic gain control (AGC) target of 300 % and maximum injection time of 246 ms. Data independent acquisition of MS^2^ spectra was performed in the Astral using loop control set to 0.7 seconds per cycle with varying isolation window widths and injection times. The following combinations were used for peptide digest: 5ms -2.5m/z, 10ms - 5m/z, 20ms - 10m/z, 40ms - 20m/z, 80ms - 40m/z. For single-cell samples: 45 ms - 15 m/z, 55ms - 15m/z, 65ms - 15m/z, 100ms - 22 m/z. A 1 m/z window overlap was used for all methods. Fragmentation of precursor ions was performed using higher energy collisional dissociation (HCD) using a normalized collision energy (NCE) of 25 %. AGC target was set to 800 %. The optimal 65ms method was used for the generated single-cell data.

### Mass spectrometry raw data analysis

All generated raw files were processed using Spectronaut version 17. Direct DIA analysis was performed in pipeline mode using default BGS factory settings unless specified. For single OCI-AML8227 cells, pulsar searches were performed without fixed modifications. Carbamidomethylation of cysteines was set as fixed modification for experiments that used diluted Hela peptides. N-terminal acetylation and methionine oxidation were set as variable modifications. Quantification level was set to MS1 and the quantity type set to area under the curve. For experiments involving the use of GPF libraries, runs were added to directDIA to supplement the single-cell dataset search.

### Single-cell analysis for cell cycle exploration

Protein quantification tables were exported from the Spectronaut software and further analyzed using custom scripts. First, quality control was applied to ensure that only cells with sufficient quality proteome profiles were used. Quality of the proteomes was determined by each cell’s total signal intensity and number of proteins measured, resulting in the filtering out of 26% of the cells (Figure S2). This left 97 cells that could be used for further analysis. Normalization was carried out in two steps: first the data was standardized sample-wise to reduce differences between total cell intensity, followed by a protein-centric standardization to remove biases induced from highly abundant proteins likely to be derived from contaminants (keratins, etc.). Missing value imputation was then carried out with K-nearest neighbor (KNN) algorithm (30% missing value cut-off). The normalized and imputed data was then used for further analysis. Principal component and uniform manifold analysis was carried out in python with the use of sclera and UMAP packages. The first two principal component values were converted to degree values with the np.arctan2 function. All data visualization was carried out in R with the use of ggplot data visualization package^41^.

### Single-cell analysis of the OCI-AML8227 model system

Bioinformatic analysis of the primary leukemia model was carried out in a protein-centric manner. For this, protein quantification tables obtained from the Spectronaut software were used to conduct the analysis. Differential expression (DE) analysis was conducted with the limma package^42^ using proteins with at least 80 % of non-missing values across all populations. Leftover missing values were imputed using the KNN algorithm. For PCA and UMAP projections, double Z-score normalization was used whereas for DE analysis, quantile normalization was used. All data analysis was carried out using R (v. 4.2.3) with ggplot as primary visualization package^41^.

## Author Contributions

V.P., P.A.F., and E.S. designed the study. V.P., P.A.F., and N.Ü. performed the experiments. V.P., P.A.F.. and B.F. performed data analysis. V.P., P.A.F., B.F., T.N.A., E.Da., and E.M.S designed the methods and evaluated the data, with input from H.S., E.De., J.P., A. P., C.H, A.M., V.Z., and B.T.P. The manuscript was drafted and revised by V.P., P.A.F., and E.S., and has been read and approved by all authors. E.M.S. supervised the work.

## Acknowledgements

This work was funded by the following grants to E.M.S.: 1) reference number NNF21OC0071016 from the Novo Nordisk Foundation; 2) case no. 2067-00053B from the Independent Research Fund Denmark. B.F. is the recipient of a fellowship from the Novo Nordisk Foundation as part of the Copenhagen Bioscience PhD. Programme, supported through grant NNF19SA0035442. V.P. is funded by a Leo Foundation grant (LF-OC-21-000832). P.A.F. is funded by a Danish Cancer Society grant (R324-A17978). Work in the Porse lab was supported by grants from the Svend Andersen Foundation, the Candys foundation, the Danish Cancer Society, and the Independent Research Fund Denmark. We also thank all members from the Cell Diversity Lab, headed by E.M.S. for constructive input and fruitful discussions.

