## Supplementary Table for "Evaluating the capabilities of the Astral mass analyzer for single-cell proteomics"

TFs

| Abundance.mean | Entry | Rank |
|---|---|---|
| 11.69309969816230 | Q8N7M2 | 50 |
| 10.113218841360000 | Q96NB3 | 173 |
| 9.10744419296398 | Q9UQ80 | 356 |
| 8.972997708737430 | P40763 | 399 |
| 8.857981693688030 | P22392 | 431 |
| 8.616322746360650 | Q9UJU3 | 502 |
| 8.521715157301370 | A6NHT5 | 544 |
| 8.189692895244820 | Q96NZ1 | 681 |
| 7.903864264234140 | Q8IWI9 | 823 |
| 7.845236378340630 | P67809 | 849 |
| 7.735017377869200 | P16989 | 899 |
| 7.678508814711910 | Q06730 | 931 |
| 7.625035340466600 | Q96LU7 | 965 |
| 7.606760075047980 | P17096 | 968 |
| 7.577351723721830 | Q8NCA9 | 985 |
| 7.570188963916120 | Q8IVH2 | 988 |
| 7.433695042768390 | P02788 | 1074 |
| 7.416517235868590 | Q9NX63 | 1088 |
| 7.278361028179470 | O75820 | 1162 |
| 7.150304167105570 | O60765 | 1243 |
| 7.1211612822346200 | P41970 | 1253 |
| 7.120254001900170 | Q9NW07 | 1254 |
| 7.102054646865300 | P78347 | 1265 |
| 7.019921611540940 | O15178 | 1322 |
| 7.007557568440320 | Q9UMN6 | 1328 |
| 6.883940605702970 | Q00577 | 1423 |
| 6.879215225853910 | P17019 | 1429 |
| 6.842636286377960 | Q9P243 | 1457 |
| 6.700478258459710 | P42229 | 1557 |
| 6.68254906371527 | Q6NUN9 | 1572 |
| 6.678312646906540 | P48382 | 1577 |
| 6.654081941855010 | P42224 | 1596 |
| 6.539910636747940 | Q8N2I2 | 1674 |
| 6.504532455159660 | Q8WYP5 | 1699 |
| 6.407884582079550 | Q14765 | 1778 |
| 6.386948747097060 | P57071 | 1795 |
| 6.376810752183750 | P17020 | 1804 |
| 6.348348915487720 | Q14526 | 1823 |
| 6.240039357697580 | P52926 | 1891 |
| 6.239601731369380 | Q86UP3 | 1892 |
| 6.189151065697230 | P0DKX0 | 1919 |
| 6.177625198285810 | Q9UHF7 | 1927 |
| 6.083764518437700 | P10072 | 1985 |
| 6.074934681482260 | Q9UFW8 | 2000 |

Epigenetic regulators

| Abundance.mean | Entry | Rank |
|---|---|---|
| 15.152112629291600 | P62805 | 12 |
| 13.967688126484800 | P10412 | 17 |
| 13.42703948851030 | P16401 | 20 |
| 11.550843130710200 | P16403 | 58 |
| 11.101350923673000 | P11142 | 80 |
| 10.670532470557200 | P06748 | 108 |
| 9.22221323918661 | Q92688 | 329 |
| 8.662668097888640 | P49321 | 491 |
| 8.585772188523890 | Q9H9B1 | 519 |
| 8.473727078414890 | Q99497 | 568 |
| 8.42520592741295 | Q13263 | 590 |
| 8.293462472286650 | O75367 | 645 |
| 8.28100197966051 | Q16576 | 648 |
| 8.25569325528907 | Q9NWY4 | 657 |
| 8.137789842920310 | Q09028 | 701 |
| 7.9699375496256600 | P49736 | 791 |
| 7.906249649454170 | Q9Y230 | 822 |
| 7.673135272312450 | Q08945 | 934 |
| 7.64922332191548 | Q9Y5B9 | 950 |
| 7.500851778931760 | Q9Y265 | 1035 |
| 7.155403219635810 | P43246 | 1242 |
| 6.979495925315100 | P55209 | 1350 |
| 6.953610583346650 | Q9NRZ9 | 1366 |
| 6.942816801720300 | Q9BTT0 | 1376 |
| 6.938028998793440 | P09874 | 1383 |
| 6.748791821456870 | O60264 | 1516 |
| 6.679862507919940 | Q99733 | 1574 |
| 6.594623026366390 | O95816 | 1641 |
| 6.550908577588680 | Q9HAF1 | 1667 |
| 6.505523548314810 | Q9NRD5 | 1697 |
| 6.498528154532420 | Q86VD1 | 1705 |
| 6.475166616000340 | P63279 | 1718 |
| 6.219305191769390 | Q7KZ85 | 1902 |
| 6.168615144580900 | O94776 | 1935 |
| 6.130309639332540 | Q9H0M4 | 1958 |
| 6.074934681482260 | Q9UFW8 | 2000 |
| 6.046310496362250 | P63165 | 2024 |
| 6.017050846155530 | P49711 | 2054 |
| 5.881580409776580 | P51532 | 2161 |
| 5.758614585020320 | P26358 | 2243 |
| 5.71036445606513 | Q92793 | 2271 |
| 5.557830082591890 | Q93009 | 2367 |
| 5.5560412452077000 | Q15020 | 2368 |
| 5.455314023938430 | Q96T88 | 2429 |

kinases

| Abundance.mean | Entry | Rank |
|---|---|---|
| 12.683574958137400 | Q8TDC3 | 32 |
| 10.438752946178400 | Q9H1R3 | 132 |
| 9.126595881556610 | O94806 | 350 |
| 9.075187511019300 | Q13315 | 363 |
| 8.42520592741295 | Q13263 | 590 |
| 8.163055957581550 | P28482 | 692 |
| 7.973366229521750 | P06493 | 788 |
| 7.870286670177320 | Q6P3W7 | 835 |
| 7.640806813952110 | P31751 | 955 |
| 7.544353190451560 | P37173 | 1001 |
| 7.449217378798380 | P78527 | 1062 |
| 7.434416100639990 | Q8TAS1 | 1072 |
| 7.423569383330790 | Q13177 | 1082 |
| 7.309626981219370 | Q05655 | 1145 |
| 7.169493539960860 | P43405 | 1232 |
| 7.1065533561826 | P16591 | 1262 |
| 7.097332500973860 | Q8WZ42 | 1275 |
| 7.064069940023450 | P15735 | 1297 |
| 6.969290029955810 | Q6ZWH5 | 1358 |
| 6.851334774518030 | Q13043 | 1450 |
| 6.819683133044420 | Q13557 | 1465 |
| 6.798267664772550 | P41240 | 1473 |
| 6.731759026565420 | P09769 | 1527 |
| 6.683404089177920 | P20794 | 1571 |
| 6.55581233914452 | P08631 | 1664 |
| 6.504628833479000 | P51812 | 1698 |
| 6.501911460841040 | Q9P289 | 1701 |
| 6.421243186854510 | P07948 | 1767 |
| 6.315279797671210 | P27361 | 1844 |
| 6.293587594300040 | O60229 | 1857 |
| 6.277826732379500 | Q16539 | 1872 |
| 6.276631321353640 | P51957 | 1873 |
| 6.263270683891350 | P53004 | 1882 |
| 6.261682564572740 | P25098 | 1884 |
| 6.138433075993350 | Q9P0L2 | 1950 |
| 6.125956988687880 | Q16644 | 1963 |
| 6.102730964978720 | Q96RR4 | 1975 |
| 6.01123656357136 | Q99986 | 2058 |
| 5.926416360068530 | O94804 | 2129 |
| 5.908287268814240 | Q00534 | 2139 |
| 5.90127370505444 | Q9NSY1 | 2145 |
| 5.794234857584370 | Q13523 | 2215 |
| 5.789306750762630 | Q15418 | 2216 |
| 5.763845009404730 | O95747 | 2235 |

| | | |
|---|---|---|
| **6.044891281600070** | Q7Z4V0 | 2026 |
| **6.02936586748361** | O15162 | 2040 |
| **6.017050846155530** | P49711 | 2054 |
| **5.989551858411380** | Q8WYA1 | 2073 |
| **5.980412534065320** | Q9NRY4 | 2085 |
| **5.960937597069230** | Q3B8N5 | 2101 |
| **5.893552790912200** | Q99459 | 2151 |
| **5.857936872007250** | Q9NQZ8 | 2172 |
| **5.85431739643497** | O43670 | 2176 |
| **5.770454430663730** | P49715 | 2229 |
| **5.758614585020320** | P26358 | 2243 |
| **5.728724380862880** | P17676 | 2256 |
| **5.70106218520498** | Q9P016 | 2278 |
| **5.65620124832786** | P35680 | 2296 |
| **5.5723449664533800** | Q13106 | 2359 |
| **5.557885568247100** | Q15424 | 2366 |
| **5.495769290930730** | Q15744 | 2401 |
| **5.489939721812110** | P18583 | 2406 |
| **5.459927083678520** | Q15075 | 2421 |
| **5.452893736763530** | O00470 | 2432 |
| **5.4043226913167900** | O95409 | 2465 |
| **5.350704821736970** | Q03701 | 2507 |
| **5.300913498024250** | Q13422 | 2543 |
| **5.233400400582230** | Q00653 | 2596 |
| **5.233303325869580** | P17483 | 2597 |
| **5.225396794990030** | Q96T58 | 2603 |
| **5.177225167024540** | Q5BKZ1 | 2632 |
| **5.093738949323510** | P49716 | 2677 |
| **5.0418222836594700** | Q01826 | 2717 |
| **4.94923077891315** | Q96JM3 | 2774 |
| **4.93673243627381** | Q96MR9 | 2787 |
| **4.826453897743150** | Q8TF47 | 2849 |
| **4.750745516851240** | Q86YP4 | 2910 |
| **4.60979438731855** | P17535 | 3004 |
| **4.585124604371320** | Q8NFU7 | 3019 |
| **4.450491992203530** | Q05516 | 3071 |
| **4.419275151032720** | Q06546 | 3085 |
| **4.249037564754470** | Q96I27 | 3152 |
| **4.227077229886060** | O60315 | 3160 |
| **4.18184116369181** | Q9NR48 | 3177 |
| **4.151173068415290** | Q14186 | 3194 |
| **4.076971989302550** | Q9HCU5 | 3220 |
| **3.9551967849711700** | Q8WXI9 | 3261 |
| **3.842464338027260** | Q49A26 | 3296 |
| **3.7552245615216700** | P35712 | 3332 |
| **3.7250206101266300** | Q15916 | 3344 |

| | | |
|---|---|---|
| **5.330201708054270** | Q7Z4V5 | 2524 |
| **5.26104002027533** | Q9BW71 | 2570 |
| **5.099266537254710** | Q14683 | 2675 |
| **5.010598101494620** | Q13112 | 2733 |
| **4.76187650769361** | Q2KHR3 | 2901 |
| **4.750745516851240** | Q86YP4 | 2910 |
| **4.66466140275797** | Q96ST3 | 2963 |
| **4.662671200231680** | Q9Y4A5 | 2965 |
| **4.585124604371320** | Q8NFU7 | 3019 |
| **4.400648146872080** | Q15022 | 3092 |
| **4.3434423395457** | Q8WXX5 | 3110 |
| **4.302619763712740** | P24941 | 3121 |
| **3.911102708576820** | Q9Y4E8 | 3278 |
| **3.8251609943706100** | P50750 | 3307 |
| **3.748208393232960** | O14744 | 3336 |
| **3.6671767966978100** | O15047 | 3360 |
| **3.661735092946060** | O60341 | 3362 |
| **2.845229441687710** | O75376 | 3507 |
| **2.71157366467734** | P40692 | 3526 |
| **2.1280145192589600** | O94874 | 3578 |
| **1.186922140230360** | P29034 | 3650 |
| **0.7697938794922350** | Q92922 | 3679 |
| **0.7424796835560210** | Q6DT37 | 3684 |
| **0.0812405854533158[illegible]** | Q6VMQ6 | 3780 |

| | | |
|---|---|---|
| **5.750345112192190** | P68400 | 2247 |
| **5.686006860871340** | Q9Y616 | 2285 |
| **5.549750691565270** | Q9H2K8 | 2374 |
| **5.479764286619520** | Q14289 | 2409 |
| **5.456812485621880** | Q9H2G2 | 2426 |
| **5.425551629436770** | Q13418 | 2452 |
| **5.416427912557160** | Q00535 | 2459 |
| **5.297934345576690** | Q9H4A3 | 2546 |
| **5.2440538089616900** | P20594 | 2586 |
| **5.2399597297229800** | P04049 | 2592 |
| **5.212154860352350** | Q8TD19 | 2610 |
| **5.196919480356280** | P53350 | 2619 |
| **5.022304899430590** | P49137 | 2728 |
| **4.987600314565030** | Q58A45 | 2744 |
| **4.791793461242890** | O95835 | 2880 |
| **4.725459654149950** | Q9Y5S2 | 2929 |
| **4.671141272493260** | Q9NRL2 | 2958 |
| **4.662671200231680** | Q9Y4A5 | 2965 |
| **4.62918845494562** | Q13153 | 2990 |
| **4.568119310535550** | Q9Y5P4 | 3030 |
| **4.494467468872870** | P29317 | 3056 |
| **4.470464466176330** | Q9UIG0 | 3066 |
| **4.444296330973690** | P19525 | 3075 |
| **4.402797620126940** | Q96S44 | 3091 |
| **4.302619763712740** | P24941 | 3121 |
| **4.195168481072610** | P42345 | 3170 |
| **4.188774359151640** | Q8TD08 | 3172 |
| **4.169002281195840** | P31152 | 3182 |
| **4.1486578900698800** | P04626 | 3197 |
| **4.00693969960142** | Q9UHY1 | 3244 |
| **3.960980418899300** | Q13464 | 3257 |
| **3.88735359860155** | O75582 | 3285 |
| **3.8269255171970100** | Q16512 | 3305 |
| **3.8251609943706100** | P50750 | 3307 |
| **3.7608179602242300** | P49840 | 3328 |
| **3.7277885259398300** | P07332 | 3343 |
| **3.4272277280722200** | Q9NSY0 | 3420 |
| **3.301545418248540** | Q7Z695 | 3441 |
| **3.266540924951200** | Q13131 | 3443 |
| **2.991261895715320** | P14616 | 3490 |
| **2.9586134069183200** | Q12979 | 3495 |
| **2.7646499894253300** | Q96J92 | 3517 |
| **2.7599990360306300** | Q13882 | 3518 |
| **2.757932559077890** | Q14296 | 3520 |
| **2.609384523644810** | Q56UN5 | 3537 |
| **2.387902348900010** | Q06187 | 3555 |

| | | |
|---|---|---|
| **3.686646318557130** | Q8TAW3 | 3355 |
| **3.4660223091285500** | Q9UJL9 | 3407 |
| **3.3594854654558200** | Q01196 | 3431 |
| **3.3560633298716700** | O15226 | 3432 |
| **3.2447937566069600** | P61244 | 3448 |
| **3.023016134484450** | P85037 | 3483 |
| **2.673871488757640** | Q9Y6Q9 | 3531 |
| **2.336006409368920** | Q13398 | 3560 |
| **2.2795662347987000** | Q9BV97 | 3564 |
| **2.2653104874122600** | P17024 | 3565 |
| **2.0777674047196800** | Q12830 | 3581 |
| **2.0199256645894700** | Q6ECI4 | 3585 |
| **1.9724963766810700** | Q9UL36 | 3588 |
| **1.8950370057400900** | P52630 | 3596 |
| **1.627544638861650** | Q14541 | 3621 |
| **1.4228418662803400** | Q9UJV8 | 3638 |
| **0.7670623976735660** | Q6ZR52 | 3681 |
| **0.7202116874438160** | Q8IYB9 | 3688 |
| **0.1356783602566810** | Q6DD87 | 3775 |

| | | |
|---|---|---|
| **2.356833184838980** | P36507 | 3557 |
| **1.5229534141783600** | Q9Y6R4 | 3631 |
| **0.7976080635344230** | Q9Y6E0 | 3673 |
| **0.7424796835560210** | Q6DT37 | 3684 |
| **0.6328936640238320** | P10721 | 3696 |
| **0.4894046515907280** | Q6ZN16 | 3718 |
| **0.2510170905843880** | Q9NYL2 | 3759 |
