## Supplementary Figures for "Evaluating the capabilities of the Astral mass analyzer for single-cell proteomics"

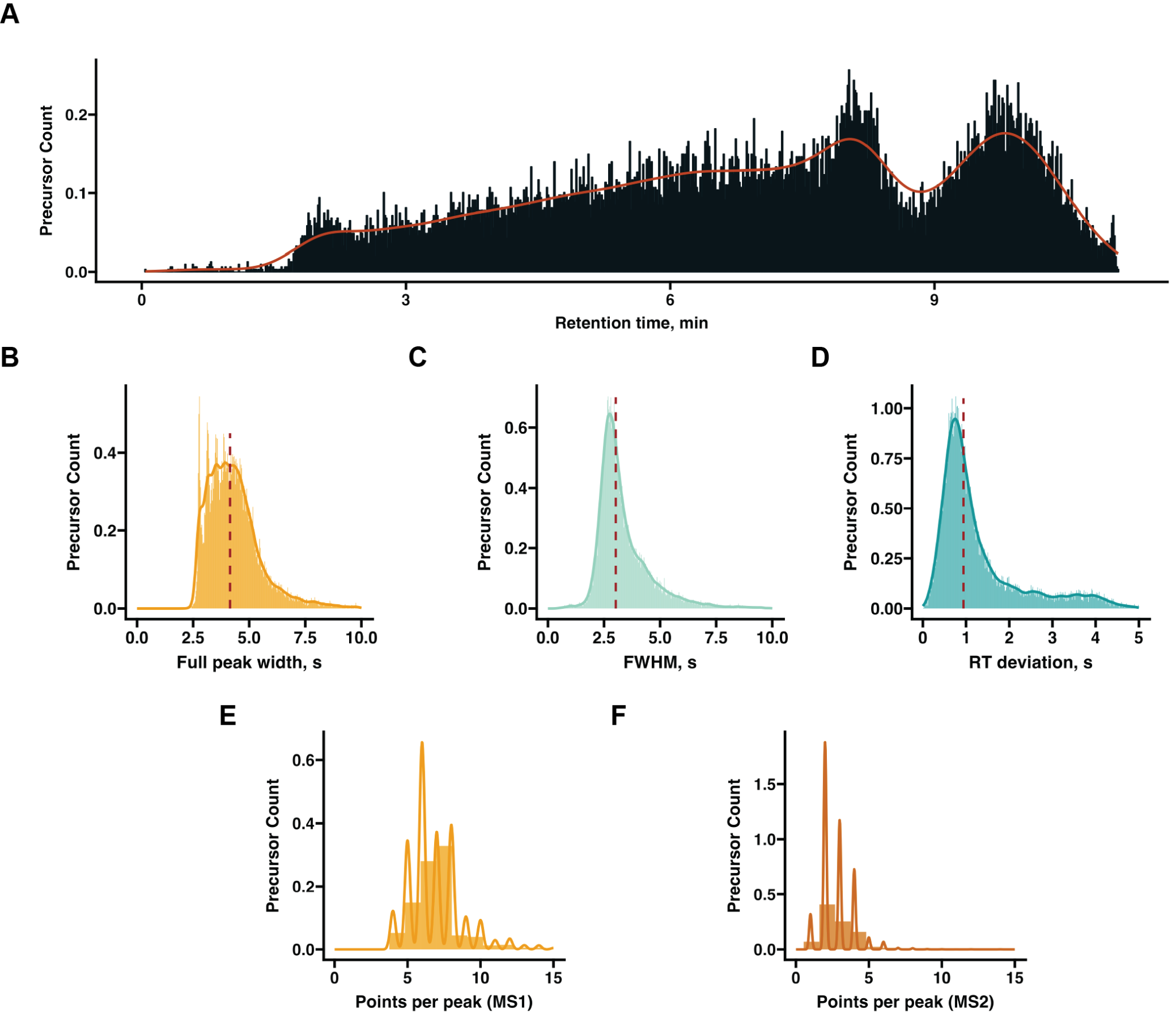

**Figure S1. Chromatographic features of the chosen chromatographic method. A)** Histogram showing precursor distribution along chosen 80SPD LC method. Histograms showing different precursor peak related properties: **B)** full-peak width, **C)** Full-width at half maximum (FWHM), **D)** Apex retention time (RT) standard deviation, **E)** Data points per peak collected on MS1, or **F)** MS2 level.

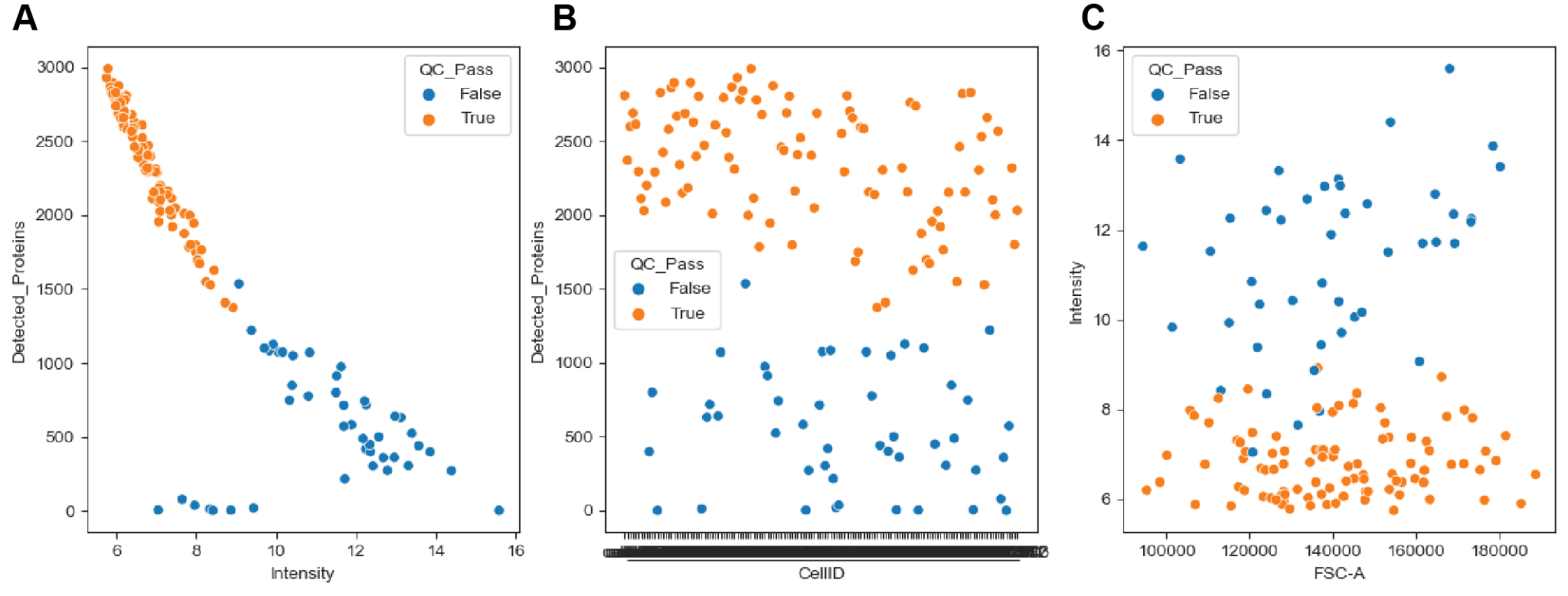

**Figure S2. Quality control of single-cell HEK293 data.** Scatter plot of quantified protein number per cell (y-axis) plotted against **A)** log transformed total sample intensities **B)** run time left to right **C)** cell are based on the forward-side scatter (FSC-A).
